# High throughput identification of genes conferring resistance or sensitivity to toxic effectors delivered by the type VI secretion system

**DOI:** 10.1101/2021.10.06.463450

**Authors:** Steven J. Hersch, Rehnuma Tabassum Sejuty, Kevin Manera, Tao G. Dong

## Abstract

The type six secretion system (T6SS) is a prevalent bacterial weapon delivering toxic effector proteins into nearby competitors. In addition to immunity genes that protect against a particular effector, alternate yet crucial nonspecific defences have also recently been identified. To systematically identify genes influencing T6SS susceptibility in numerous species, we designed a Tn-Seq-based competition assay. Combined with follow-up analyses using *E. coli* and *V. cholerae* gene knockout collections, we demonstrate that our Tn-Seq competition technique can be used to identify both immunity and non-immunity defences against the T6SS. We also identify *E. coli* proteins that facilitate T6SS-mediated cell death, including metabolic genes such as *cyaA* and *gltA,* where mutant strains were resistant to attack. Our findings act as a proof-of-concept for the technique while also illuminating novel genes of interest. Since Tn-Seq can be applied in numerous species, our method has broad potential for identifying diverse T6SS defence genes across genomes in a high-throughput manner.

**Importance:** The type six secretion system (T6SS) is a molecular poison-tipped spear that bacteria use to kill nearby competitors. To prevent self killing, they use antitoxins called immunity genes that specifically neutralize the poisons. Beyond immunity genes, multiple additional defences have recently been discovered but there are likely many more across the genomes of diverse species. To help discover these novel mechanisms, we designed a high-throughput method that can be used in numerous different species to rapidly identify genes involved in sensitivity to T6SS attacks. Using T6SS ‘killers’ delivering individual poisons and two commonly studied ‘prey’ bacteria, we show proof-of-principle that the technique can discover proteins that make the prey cells more resistant or sensitive to particular poisons. This will greatly improve the speed at which genes influencing the T6SS can be identified and selected for further study in follow-up analyses.

## Introduction

Microbes are constantly competing for limited space and nutrients and have evolved many mechanisms for outcompeting their neighbours. These adaptations include weapons used to kill or inhibit competitors, such as the bacterial type six secretion system (T6SS). Resembling a molecular spear, the T6SS is employed by approximately 25% of Gram-negative bacteria to stab nearby cells and deliver toxic ‘effector’ proteins mounted on the T6SS structure (1–4). Importantly, bacteria with antibacterial T6SS effectors also encode immunity genes to protect themselves against their own weapon.

While immunity genes remain the gold standard of defence against T6SS effectors, numerous non-immunity defences have also been discovered (5, 6): These include stress responses that help cells survive damage caused by effectors(7–10), physical separation mechanisms (11–13), exopolysaccharides that act as armour to deflect incoming T6SS attacks (7, 14, 15), cell wall modifying enzymes that render the target molecules resistant to effectors (16), and proteins that play a role in survival by unclear mechanisms such as the periplasmic chaperone, Spy, the periplasmic protease inhibitor, Ecotin, and the outer membrane maltose porin, LamB, amongst others (7, 17, 18). Prey cell proteins can also play a role in activating incoming effectors: The ClpAP protease complex was shown to enhance susceptibility to the *A. tumefaciens* T6SS (19), and the *Serratia marcescens* effectors, Ssp2 and Ssp4, get activated in prey cells when their DsbA protein generates an intramolecular disulfide bridge (20). Notably, periplasmic disulfide bridges formed by DsbA are also important for the functionality of numerous immunity genes (21–25). The metabolic state of the prey cell can also have a profound influence on survival: Stationary phase prey can be more susceptible to attacks due to their decreased ability to form microcolonies isolated from attackers, replace damaged components or produce sufficient immunity proteins for protection (7, 11, 12, 26–28). Finally, catabolite repression is another mechanism recently shown to influence *E. coli* survival against a T6SS attack, where growth with glucose or using mutant prey lacking the cAMP receptor protein (CRP) showed improved survival against the T6SS of *Vibrio cholerae* strain C6706 (29).

*V. cholerae* strain V52 encodes a well-studied T6SS and four antibacterial effectors: VgrG3 and TseH both target peptidoglycan, TseL targets lipids, and VasX encodes a colicin-like domain (30–33). A fifth effector, VgrG1 encodes an actin cross-linking domain (34). Notably, while each of the antibacterial effectors has an associated immunity gene, prey in stationary phase or mutants lacking the cell wall damage-sensing two-component system, VxrAB (WigKR), become susceptible to killing by T6SS-active sister cells (7). These findings highlight that there are numerous mechanisms for defending against T6SS effectors and some of them are required for full survival even in the presence of immunity genes.

While multiple non-immunity factors influencing T6SS effector-susceptibility have now been characterized, most have been identified in a hypothesis-driven manner or using time-intensive screens of knockout libraries. These techniques have worked well as proof-of-principle for these defence systems, but leave much of the bacterial genome unexplored for influencing T6SS activity. Transposon insertion sequencing (Tn-Seq) is a method where a transposon is randomly inserted into the bacterial genome to generate a large mutant library with single insertions across the genome (35–38). Using next-generation sequencing, the location and abundance of each insertion site can be quantified in parallel and compared between control and test conditions. If a transposon impairs a gene crucial to survival, that insertion site will be depleted in the test condition relative to the control. Tn-Seq is a high-throughput method that can be employed in many different species and probe the majority of the genome in parallel.

In this work, we used Tn-Seq transposon insertion libraries as prey in competition with T6SS^+^ killer strains delivering individual effectors. Using this method, we successfully identify numerous genes in both *E. coli* and *V. cholerae* prey species that play significant roles in rendering the prey susceptible or resistant to specific T6SS effectors. This technique can be readily adapted to diverse prey species and killer strains for high-throughput, systematic exploration of genes influencing effector specificity and T6SS susceptibility.

## Results

### Tn-Seq reveals *E. coli* mutants that are resistant to the *V. cholerae* T6SS effector, VgrG3

To test the ability of a Tn-Seq competition assay to identify genes influencing survival against T6SS effectors, we first examined *E. coli* prey. *E. coli* are commonly used as prey cells to measure the activity of T6SSs and we previously examined how specific gene knockouts can drastically reduce *E. coli* survival against the *V. cholerae* T6SS effectors, TseH and TseL (7, 8). Other groups have also identified genes that influence *E. coli* survival against T6SSs, including mutants in *clpP* or *crp* genes, which were resistant to the T6SS of *Agrobacterium tumefaciens* or *V. cholerae*, respectively (19, 29). Cumulatively, these works provide a set of genes with known roles in T6SS survival, which serve as positive controls for identifying important genes. Additionally, a genome-wide single gene knockout collection exists in *E. coli* (the Keio collection) for secondary screening of genes of interest identified by the Tn-Seq competition assays (39).

In our lab conditions, we previously found that the *V. cholerae* effectors, TseL and TseH, do not show significant killing of wildtype *E. coli* prey (7, 8). Using a high killer:prey ratio, we hoped to identify genes required for survival when challenged by these effectors. We generated an *E. coli* transposon insertion library and compared Tn-Seq reads recovered after incubation with T6SS-null control or T6SS-active killer cells delivering individual effectors. For incubation with killers delivering TseL or TseH, we used a fold-change (T6SS-null/active) cutoff of 10 and a p-value cutoff of 0.05. However, for these effectors, the genes identified as hits by our analysis were almost entirely pseudogenes, tRNAs, or non-coding RNAs (ncRNA) (Table S1, Figure S1). Therefore, these likely reflect a propensity for transposon insertion at these sites or artifacts of sequencing rather than true roles in survival against the T6SS effector. Because of this, we did not follow up on the results for these two effectors.

In contrast to TseL and TseH, our Tn-Seq competition analysis with *V. cholerae* delivering VgrG3 was more promising. Since VgrG3 is potent against *E. coli*, we used a 1:1 killer:prey ratio to recover sufficient prey cells following the 3-hour competition. We used the same significance thresholds and screened out the likely artifacts described above. Numerous genes were identified to be significantly reduced or more abundant in the VgrG3-treated population compared to controls, suggesting sensitivity or resistance (respectively) to the effector (Fig. 1A). Notably, the resistant mutants included the central metabolism gene encoding citrate synthase (*gltA*), the catabolite repression gene, adenylate cyclase (*cyaA*), and the *clpP* protease. Sensitive mutants identified included a peptidoglycan maturation protein, *mtgA*, and a subunit of the maltose transporter, *malG.*

**Figure 1:**
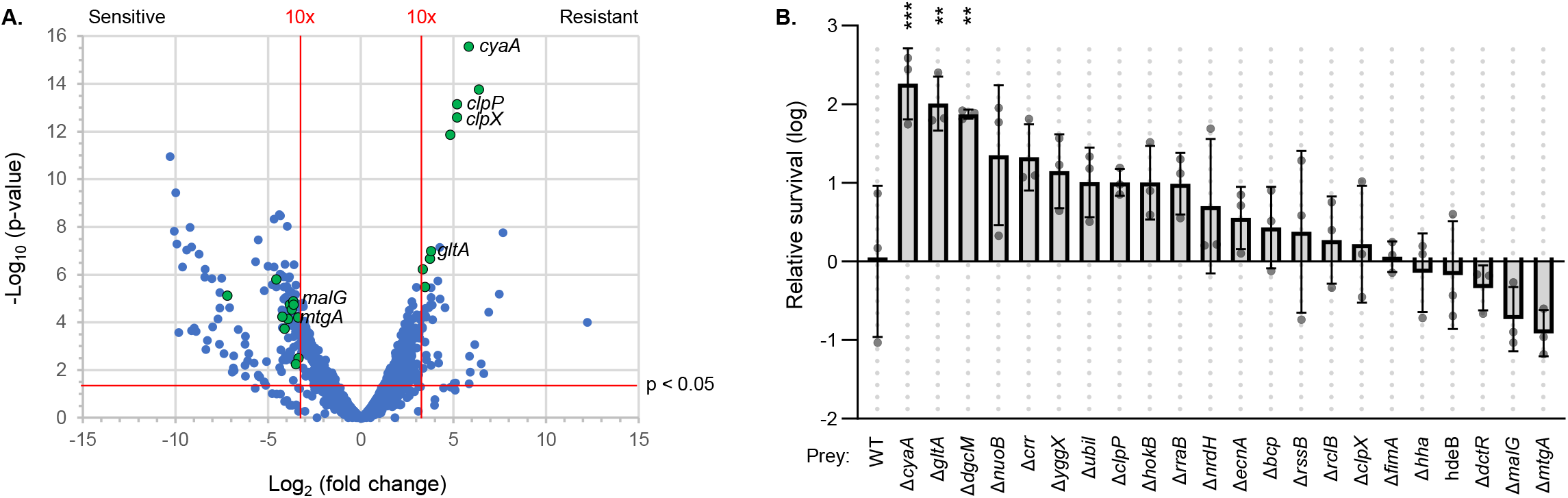
*E. coli::Tn* survival against *V. cholerae* VgrG3^+^. **A.** Volcano plot overviewing Tn-Seq data with *E. coli* prey. X-axis shows fold change comparing *V. cholerae* killer strains with only VgrG3 intact (VgrG3^+^) or all antibacterial effectors inactivated (4eff_C_). A negative number shows reduced abundance in the sample treated with VgrG3^+^.Vertical red lines indicate ~10-fold changes. Y-axis shows p-value of EDGE test statistical analysis. Horizontal red line indicates ~0.05. All data are shown as blue dots and genes selected for follow-up analysis are indicated as green dots with black outlines. Select genes of interest are labeled to the right of the corresponding datapoint. **B.** Bacterial competition assay follow-up analysis on select genes of interest. Data shows relative survival (as log) comparing CFU of *E. coli* prey after incubation with VgrG3^+^ / 4eff_C_ *V. cholerae* killer strains. This ratio was further normalized to the wild-type (WT) prey strain for clarity of comparison. One way ANOVA with Dunnett’s multiple comparisons test comparing samples to WT prey: *** p < 0.001; ** p < 0.01.

To further examine the genes identified by Tn-Seq, we employed the *E. coli* Keio collection of single gene knockout strains (39). We competed Keio collection mutants (selected based on the Tn-Seq hits) or the *E. coli* BW25113 wildtype strain with *V. cholerae* delivering VgrG3 or a T6SS-null control. While variance in the survival of wildtype *E. coli* limited the statistical power of this experiment, mutants lacking the *gltA* or *cyaA* genes showed significantly improved survival against VgrG3 (Fig. 1B). Though *clpP* and other mutants averaged over 10-fold improved survival, they did not reach a p-value below 0.05 in this experiment. Similarly, *mtgA* and *malG* averaged over 5-fold reduced survival compared to wildtype, but were not significantly different due to the variance in the wildtype control. Interestingly, another mutant, *dgcM* (*ydaM*), also showed significant resistance to VgrG3 in this assay (Fig. 1B). The *dgcM* gene encodes a diguanylate cyclase that is best characterized for its role in regulating curli biosynthesis (40).

### *cyaA* and *dgcM* mutants are also resistant to the *V. cholerae* T6SS effector, VasX

Similar to VgrG3, *E. coli* prey are also susceptible to killing by the *V. cholerae* T6SS effector, VasX. We again employed Tn-Seq competition to assay mutants that are sensitive or resistant to this effector. Using the same assay parameters as for VgrG3, we found a small set of genes with significantly reduced or increased abundance in the VasX-treated sample compared to the T6SS-null control (Fig. 2A). The reactive chlorine stress protein, *rclB*, and the H-NS-interacting protein, Hha, showed significant sensitivity to VasX in the Tn-Seq assay. However, they did not show significantly reduced survival in the follow-up analysis using the equivalent Keio collection knockouts (Fig. 2B). Interestingly, *cyaA* and *dgcM* mutants showed significantly improved survival against VasX in both the Tn-Seq competition and the follow-up experiment (Fig. 2), demonstrating that these mutant strains are resistant to VasX in addition to VgrG3.

**Figure 2:**
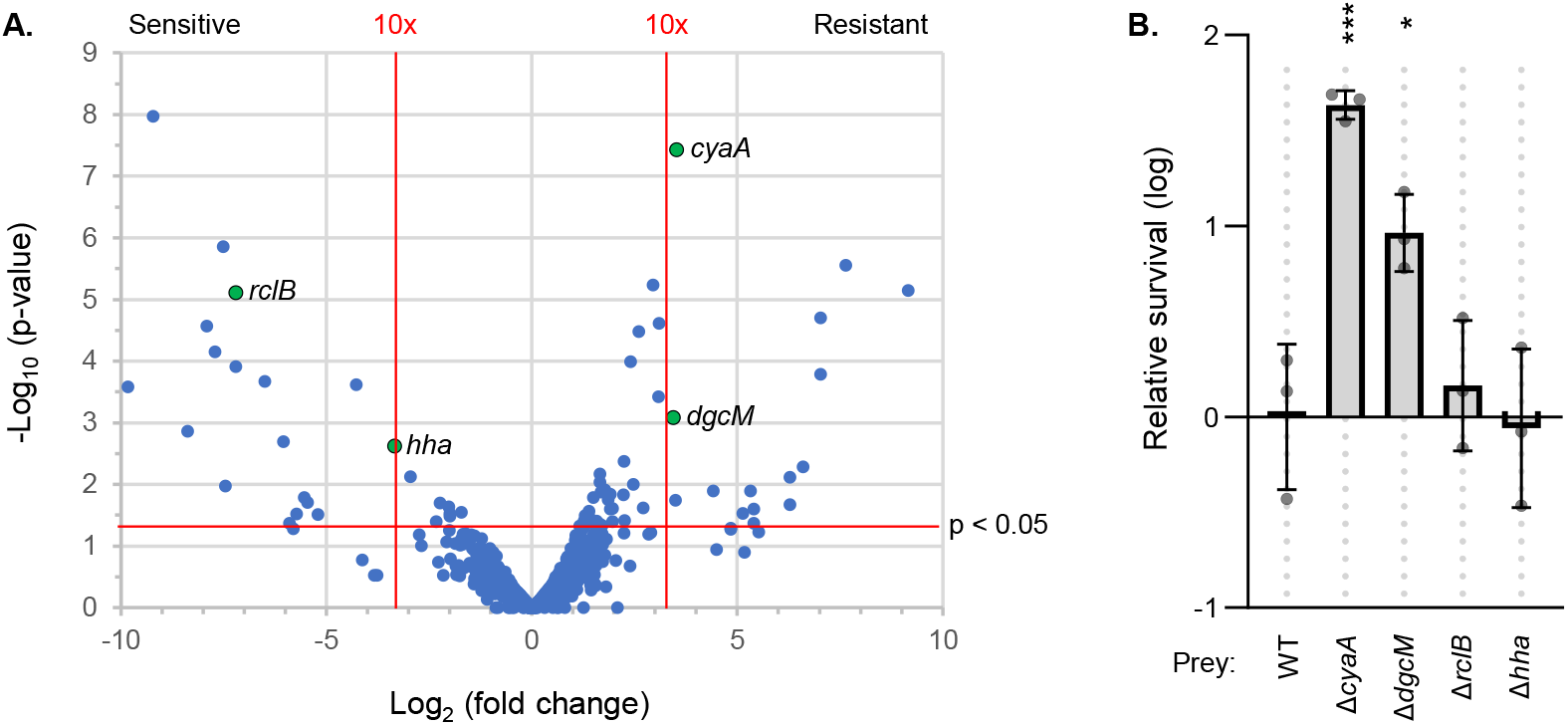
*E. coli::Tn* survival against *V. cholerae* VasX^+^. **A.** Volcano plot overviewing Tn-Seq data with *E. coli* prey. X-axis shows fold change comparing *V. cholerae* killer strains with only VasX intact (VasX^+^) or all antibacterial effectors inactivated (4eff_C_). A negative number shows reduced abundance in the sample treated with VasX^+^.Vertical red lines indicate ~10-fold changes. Y-axis shows p-value of EDGE test statistical analysis. Horizontal red line indicates ~0.05. All data are shown as blue dots and genes selected for follow-up analysis are indicated as green dots with black outlines. Select genes of interest are labeled to the right of the corresponding datapoint. **B.** Bacterial competition assay follow-up analysis on select genes of interest. Data shows relative survival (as log) comparing CFU of *E. coli* prey after incubation with VasX^+^ / 4eff_C_ *V. cholerae* killer strains. This ratio was further normalized to the wild-type (WT) prey strain for clarity of comparison. One way ANOVA with Dunnett’s multiple comparisons test comparing samples to WT prey: *** p < 0.001; * p < 0.05.

### Tn-Seq reveals *V. cholerae* mutants that are sensitive to the *V. cholerae* T6SS

We previously showed that when *V. cholerae* prey are challenged by isogenic killers, the prey require the VxrAB (WigKR) two-component system for full survival despite encoding all immunity genes (7). This highlighted that additional systems may be required in parallel to immunity genes for full protection. To further explore defences required even in the presence of immunity proteins, we again used our Tn-Seq competition assay. Using a 10:1 killer:prey ratio, we competed a *V. cholerae* V52 Tn-library (prey) against wildtype or T6SS-null V52 (killers). The results of this analysis demonstrated numerous mutants that were drastically reduced in the T6SS^+^ wildtype-challenged sample compared to control. As a demonstration of the assay’s veracity, these genes included all three of the immunity genes (*tsiV1-3*) with effectors that kill *V. cholerae* (Fig. 3A; Table S1); the fourth immunity gene, *tsiH*, is not required for full survival against TseH (7, 30). The high-throughput assay identified numerous other transposon mutants with significantly reduced survival including *dsbA* (VC0034), a well-characterized oxidoreductase that catalyzes disulfide bond formation in the periplasm. Surprisingly, we also found that transposon insertion into PAAR1 (VCA0105), the first gene in the main *V. cholerae* T6SS operon, led to increased relative survival. This suggests that this transposon mutant was resistant to T6SS attacks.

**Figure 3:**
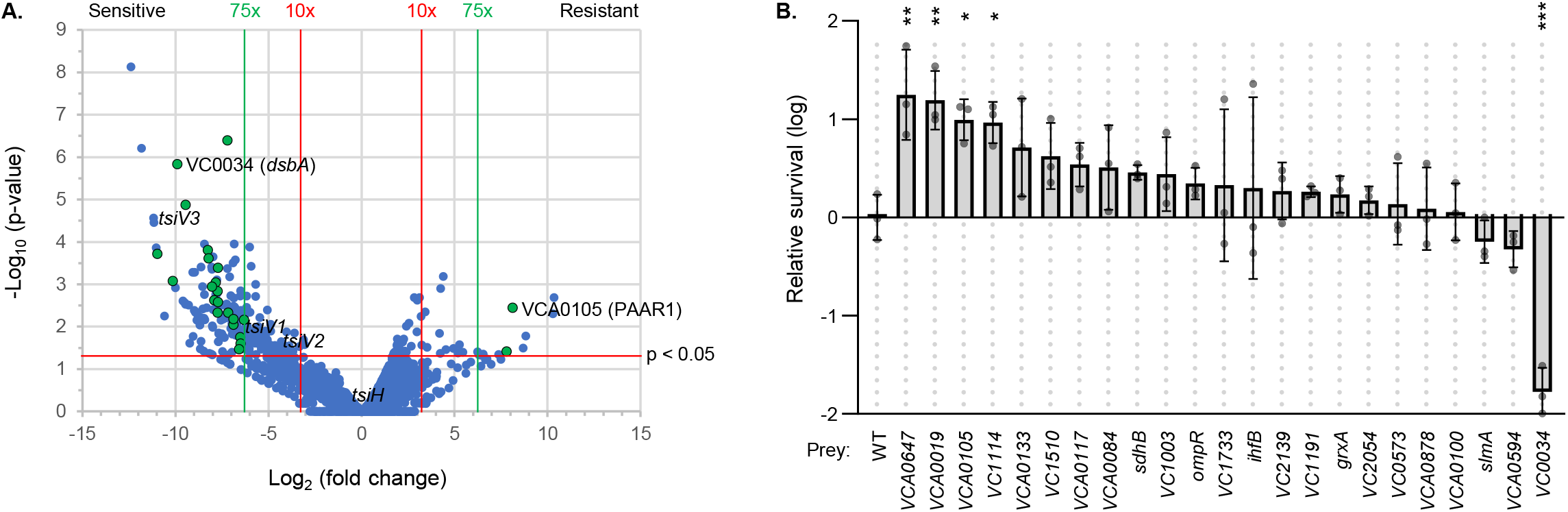
*V. cholerae::Tn* survival against *V. cholerae* T6SS. **A.**Volcano plot overviewing Tn-Seq data with *V. cholerae* prey. X-axis shows fold change comparing wild-type *V. cholerae* and Δ*TssM* killer strains (negative number shows reduced abundance in the sample treated with wild-type).Vertical red lines indicate ~10-fold changes and green lines indicate ~75-fold changes. Y-axis shows p-value of EDGE test statistical analysis. Horizontal red line indicates ~0.05. All data are shown as blue dots and genes selected for follow-up analysis are indicated as green dots with black outlines. Select genes of interest are labeled to the right of the corresponding datapoint. **B.** Bacterial competition assay follow-up analysis on select genes of interest. Data shows relative survival (as log) comparing CFU of *V. cholerae* prey after incubation with wild-type / Δ*TssM V. cholerae* killer strains. This ratio was further normalized to the wild-type (WT) prey strain for clarity of comparison. One way ANOVA with Dunnett’s multiple comparisons test comparing samples to WT prey: *** p < 0.001; ** p < 0.01; * p < 0.05.

As a follow-up screen of select genes, we employed a defined transposon mutant collection generated previously in *V. cholerae* strain C6706 (41). Using this collection, we identified four mutants that showed significantly higher T6SS resistance than wildtype C6706 (Fig. 3B). These included PAAR1, supporting the Tn-Seq competition data. Surprisingly however, the other three (VCA0119, VCA0647, VC1114) were sensitive to the T6SS attack in the Tn-Seq data. Therefore, for these genes there was a discrepancy between the two assays, possibly due to using different *V. cholerae* strains in each method. Because of this, we sought to further support the findings for PAAR1 by examining it in the V52 background using a scarless gene knockout generated by homologous recombination rather than transposon insertion. The V52 wildtype was already significantly more resistant than C6706 in this assay, making it difficult to assess if PAAR1 deletion granted further protection in this background (Fig. S2).

### *V. cholerae* requires DsbA to protect against the effector, TseL

In our follow-up analysis using C6706 mutants, the *dsbA* mutant showed greatly reduced survival compared to wildtype C6706 (Fig. 3B), supporting the Tn-Seq competition data for this gene (Fig. 3A; Table S1). We were curious if the loss of DsbA activity resulted in sensitivity to the T6SS in general or to a specific *V. cholerae* effector. To test this, we used V52 killer strains with single active effectors; all other antibacterial effectors were catalytically inactivated. The *V. cholerae* C6706 *dsbA* mutant was sensitive to killer cells delivering TseL but not other individual effectors (Fig. 4A; S3A). An *E. coli dsbA* mutant was no more sensitive to any effector (Fig. 4B; S3B). These findings suggest that DsbA helps protect against TseL in *V. cholerae*.

**Figure 4:**
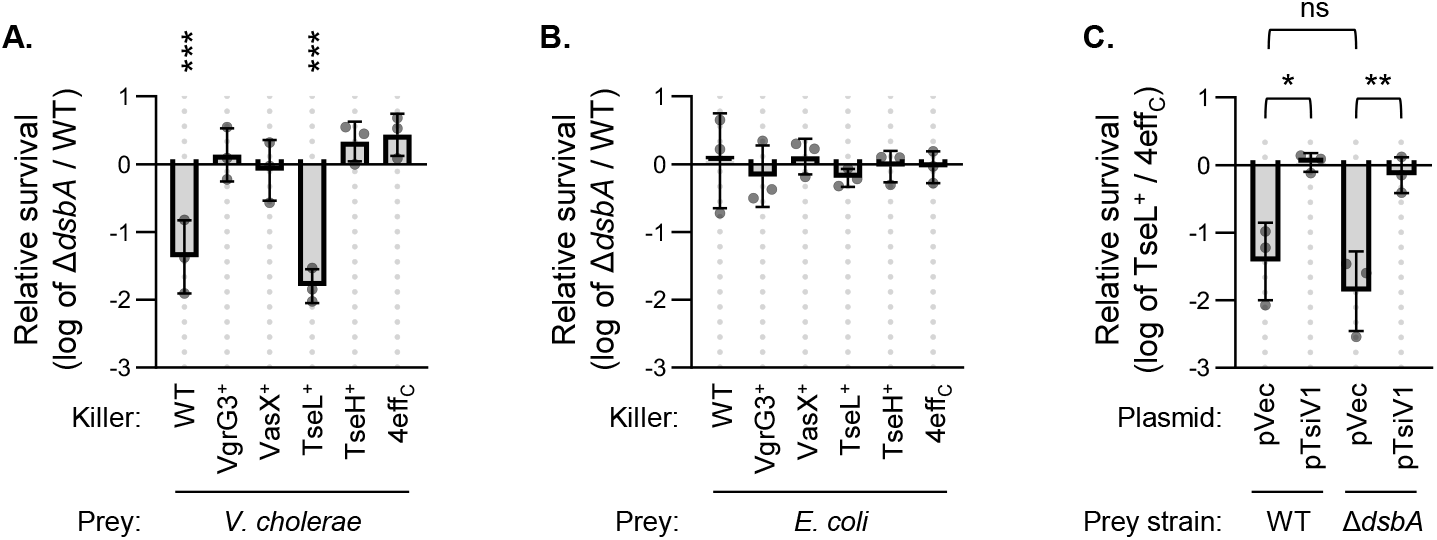
DsbA protects against TseL in *V. cholerae* but not *E. coli*. **A.** Relative survival (log of Δ*dsbA* / wild-type) of *V. cholerae* prey after competition with wild-type (WT) *V. cholerae*, a strain with all antibacterial effectors catalytically inactivated (4eff_C_), or killer strains with only one active effector (indicated). One way ANOVA with Dunnett’s multiple comparisons test comparing samples to 4eff_C_ strain: *** p < 0.001. **B.** As in A. but with *E. coli* prey. **C.** Relative survival of *E. coli* prey after competition with *V. cholerae* killers strains with only TseL active (TseL^+^) relative to all antibacterial effectors catalytically inactivated (4eff_C_). Prey *E. coli* strains were the Keio collection *dsbA* knockout or the wildtype background and contain plasmids expressing the TseL immunity gene (pTsiV1) or vector control (pVec). 0.3mM each of CaCl_2_ and MgCl_2_ were added to competition media to enhance TseL toxicity towards *E. coli*. One way ANOVA with Tukey’s multiple comparisons test: ** p < 0.01; * p < 0.05; ns, not significant.

We hypothesized that the immunity protein for TseL, TsiV1, may require a disulfide bond formed by DsbA. In this case the loss of *dsbA* would lead to impaired TsiV1 function and protection in *V. cholerae* but would have no effect in *E. coli*, which does not encode *tsiV1* and is also naturally resistant to the TseL effector (8). To test this hypothesis, we first sensitized *E. coli* to TseL: Based on differences in LB media that influence *E. coli*’s natural resistance to TseH, we added CaCl_2_ and MgCl_2_ to the competition assay media (42). Under these conditions we observed significant killing of *E. coli* by TseL, with similar toxicity to the *dsbA* mutant as wildtype (Fig. 4C). We then expressed *tsiV1* from a plasmid in *E. coli* to see if *dsbA* was required for its protective effect. Interestingly, we found that the TsiV1 plasmid was fully protective even in the *dsbA* mutant (Fig. 4C), suggesting that *dsbA* was not required for functionality of the immunity gene.

## Discussion

In this work we employed Tn-Seq high-throughput assays and a set of single-effector-active mutants to identify candidate genes influencing prey susceptibility and specificity of T6SS effectors. We demonstrated the power of this technique to decipher *E. coli* susceptibility to individual *V. cholerae* effectors, and *V. cholerae* survival against its isogenic T6SS. Cumulatively these results highlight that T6SS-mediated competition poses strong selection for multiple cellular pathways in addition to immunity genes.

Two mutants identified as more sensitive to VgrG3 in the Tn-Seq competition assay also showed reduced survival on average in our follow-up screen (Fig. 1). Though the reduction in Keio mutant survival did not reach a significant p-value, these genes may play a role in sensitivity to VgrG3. MtgA is involved in peptidoglycan maturation and so may logically support cell wall robustness for surviving challenges from VgrG3, which targets peptidoglycan. A role for the maltose transporter, MalG, is less clear, however maltose import genes have been implicated previously in survival against the *V. cholerae* V52 T6SS: Deletion of the maltose porin, LamB, was protective against V52’s T6SS and overexpression of *lamB* rendered *E. coli* more sensitive to the T6SS attack (17). How maltose transport proteins influence T6SS sensitivity remains to be determined; notably, they are regulated by catabolite repression.

Recent work examining the influence of glucose and catabolite repression demonstrated a significant role for this system in *E. coli* survival against the *V. cholerae* T6SS (29). Specifically, the authors of that work found that glucose supplementation promotes prey survival against T6SS effectors. Accordingly, the prey were similarly protected if they lacked the cAMP receptor protein, CRP, which binds to cAMP to regulate a diverse array of genes when glucose is absent. In alignment with that work, here we find that deletion of *cyaA* (encoding adenylate cyclase, which generates cAMP) is protective against the *V. cholerae* effectors, VgrG3 and VasX (Fig. 1, 2). Moreover, we show that deletion of the first gene of the tricarboxylic acid (TCA) cycle, citrate synthase (encoded by the *gltA* gene), also renders *E. coli* resistant to VgrG3 (Fig. 1). The *gltA* gene is regulated by CRP and its deletion instigates a key shift in metabolic flux. This likely leads to accumulation of acetate and acidification of the medium, which Crisan *et. al*. also tested and demonstrated to be important during glucose-mediated T6SS protection (29). In summary, our data support their work that catabolite repression and the shutdown of metabolic flux through the TCA cycle can improve *E. coli* survival against *V. cholerae* T6SS effectors. In addition, the alignment of our findings with their previous work emphasizes that our Tn-Seq competition assay can successfully identify important genes for T6SS sensitivity.

We also identified that deletion of the *dgcM* gene improved *E. coli* survival against both VgrG3 and VasX. DgcM is best known for its role in regulating the expression of curli, an amyloid fiber protein produced as a component of biofilms (40). We and others have demonstrated that biofilm components can have a drastic influence on surviving T6SS attacks, including colanic acid capsules that appear to deflect incoming T6SS spears (7, 11, 12, 14, 15). Likely, deletion of *dgcM* leads to an altered cell envelope that is more resistant to T6SS attacks. Potentially, the cells may sense the impaired curli expression in this mutant and instead increase expression of colanic acid, which subsequently protects against the T6SS (7, 14). Alternatively, since reduced curli synthesis in *dgcM* mutants leads to reduced cellular aggregation (40), it’s possible that this mutation restricts contact with attacking cells. Either mechanism could result in improved survival against multiple effectors by preventing T6SS delivery.

We tested Tn-Seq competition assays using killer strains delivering only TseL or TseH, but we did not get reliable results from those effectors. This illuminates a limitation of the method where effectors that only negligibly inhibit the prey strain may not yield sufficient killing to show reliable dropout of sensitive populations from the Tn-Seq analysis. Future studies using this method to examine modest effector activities could employ environmental conditions that increase effector efficacy, such as supplementing magnesium and calcium, which improve TseL and TseH killing (Fig. 4C) (42).

Our Tn-Seq-based method was also useful in prey encoding all immunity genes. The immunity genes of *V. cholerae* are already known in this proof-of-principle, but the method could be employed in other species to identify immunity genes and their associated effectors. A similar approach was employed previously using a T6SS-active transposon library (33). The advantage of the competition-based approach used here is that a transposon insertion that influences both T6SS expression and survival can be detected since the killer strain is not mutated. Another advantage is that the prey cells can be in a different metabolic state than the killers. Since some killing of isogenic prey is observed with stationary phase prey, this allowed us to also screen for mutants that are more resistant to the T6SS attack.

Interestingly, we found that transposon insertion in VCA0105 (PAAR1), at the beginning of the main T6SS cluster, resulted in over 200-fold resistance to attack in the Tn-Seq assay and also a significant increase in prey survival in the follow-up experiment (Fig. 3). This was in contrast to numerous other genes downstream in the T6SS cluster, which showed reduced survival in the Tn-Seq assay (Table S1). The mechanism for this is unclear. Potentially, transposon insertion mid-operon interrupts expression of the TsiV3 immunity gene (VCA0124) at the end of the operon. Indeed, Tn-insertion in TsiV3 showed 2000-fold reduced survival in the Tn-Seq competition analysis. In contrast, transposon insertion near the promoter (in PAAR1) may influence expression to potentially increase immunity gene expression in stationary phase. Supporting this hypothesis, we found that the C6706 strain (repressed T6SS under lab conditions) was significantly more sensitive to the T6SS attack than V52 (constitutively active T6SS) (Fig. S2).

Notably, we identified discrepancies between the Tn-Seq competition results and data from our follow-up assays with C6706. Specifically, we originally identified VCA0019, VCA0647, and VC1114 as sensitive to the T6SS in the Tn-Seq analysis, but then they showed significant resistance in the follow-up experiment (Fig. 3). One possibility is that the specific insertion site of the transposons in each assay may have differing effects on nearby gene expression, such as the immunity gene TsiV2 (VCA0021). Another possibility is a difference between the V52 and C6706 strain backgrounds. As noted above, wildtype V52 was significantly more resistant to the T6SS attack than C6706 was, and this difference in baseline resistance may also contribute to the difference in the sensitivity of the transposon insertion mutants. In addition, survival of mutants may also be affected when they exist in a complex mutant pool during the Tn-seq screening in comparison with the killer-prey duo in the follow-up assays.

Of the genes tested that showed increased sensitivity to T6SS attack in the Tn-Seq competition assay, only the *dsbA* mutant also showed significantly reduced survival in the follow-up screen with C6706. DsbA forms disulfide bonds in the periplasm and several T6SS effector immunity genes have been demonstrated to require *dsbA* and/or disulfide bonds to function properly (21–25). Indeed, we found that the *dsbA* mutant of *V. cholerae* – but not *E. coli* – was specifically sensitive to the TseL effector (Fig. 4, S3). However, when we expressed the *tsiV1* gene in *E. coli*, it was still protective in the *dsbA* mutant (Fig. 4C). This suggests that either a different *E. coli* gene can catalyze disulfide bond formation in TsiV1, or that DsbA influences *V. cholerae*’s sensitivity to TseL by a mechanism other than via TsiV1. Since the *E. coli dsbA* mutant was no more sensitive than wildtype, this mechanism appears to be divergent between the two tested species. While the details of this defence require further study, DsbA influences many cellular processes including envelope maintenance, which is important for surviving T6SS attacks (7, 8). Importantly, DsbA is required for colonization of *V. cholerae* in infant mice and for secretion of cholera toxin (43, 44). Our findings reveal an important and yet unrecognized role of the T6SS in selecting for the maintenance of DsbA and thus cholera toxin secretion during *V. cholerae* pathogenesis. Accordingly, bile salts can alter DsbA activity to regulate virulence gene expression (45) but have also been shown to modulate T6SS activity in pandemic strains of *V. cholerae* (46). Finally, our findings also suggest that DsbA inhibitors could render some bacteria sensitive to their own T6SS (47).

In summary, the Tn-Seq competition assay described here has great potential as a high-throughput method to identify candidate genes in numerous prey species that influence susceptibility to T6SS effectors.

## Materials and Methods

### Bacterial strains and growth conditions

Strains and plasmids used in this study are listed in Supplementary Table S2. All *V. cholerae* strain V52 were from the ‘RHH’ background with deletions of *rtxA*, *hlyA*, and *hapA* (1). *V. cholerae* C6706 mutant strains were from a defined transposon insertion collection (41). Bacteria were grown shaking at 37 °C in LB media (0.5% NaCl) or on LB agar plates. For plasmid maintenance, transposon selection, and selection of prey cells for C.F.U. counts, antibiotics were added to a final concentration of 50 μg/mL (kanamycin), 25 μg/mL (chloramphenicol), or 20 μg/mL (gentamicin).

### T6SS competition assay

Assessment of T6SS killing activity was conducted as described previously with minor modifications (7). In brief, ‘killer’ strains were subcultured 1/100 from overnight cultures for 3 hours (to an OD_600nm_ ~ 1.0 - 1.5). Prey strains were from liquid overnight cultures, with the exception of Tn-Seq prey that were resuspended from colonies grown overnight on LB agar plates (see Tn-Seq method). Killer and prey were mixed at a 10:1 ratio for *V. cholerae* prey or when using TseL^+^ or TseH^+^ killer strains. A 1:1 ratio was used for *E. coli* prey in competition with VgrG3^+^ or VasX^+^ killer strains. Mixtures were spotted on LB plates, incubated for 3 hours at 37 °C, and agar plugs containing the mixed bacteria were removed using wide-bore pipette tips, resuspended in PBS, serially diluted and plated for colony forming units (CFU). Data are reported as either log CFU recovered or as a ratio of log CFU recovered relative to a ΔT6SS control killer strain.

### Tn-Seq competition assay and analysis

The Tn-Seq competition assay was based on Tn-Seq analysis described previously (33) combined with a T6SS competition assay. In brief, the pSAMDGm suicide plasmid was conjugated into prey cells to deliver a mariner transposon with MmeI sites in the inverted repeats (33, 48). The conjugation donor strain was WM6026 (49), which requires the addition of diaminopimelic acid (DAP) to grow, allowing selection of recipients solely by the transposon resistance gene (gentamicin). Recipient cells were *E. coli* MG1655 or *V. cholerae* V52 *ΔtssM*; the deletion in *tssM* is to prevent T6SS activity in these prey cells. Transposon recipients were recovered as individual colonies (~80,000 colonies for *E. coli* libraries, ~65,000 for *V. cholerae*), then the plates were flooded with LB media to pool all colonies and generate a Tn-library.

The Tn-library was immediately used as the prey strain in a T6SS competition assay with mid-log phase *V. cholerae* killer cells. *V. cholerae* Tn-library was competed against WT or *ΔtssM* V52 killers at a 10:1 ratio. *E. coli* Tn-library was competed against VgrG3^+^, VasX^+^ or *ΔtssM* V52 killers at a 1:1 ratio, or against TseL^+^, TseH^+^ and the *ΔtssM* control at a 10:1 ratio. The mixtures of killer and prey cells were spotted onto LB agar for 3 hours as in a typical T6SS killing assay. Agar plugs containing the mixed bacteria were then removed using wide-bore pipette tips, resuspended in PBS, and distributed on enough LB plates to acquire individual colonies (~20,000 total colonies per replicate). These ‘survivor’ colonies were then pooled and prepped for Tn-Seq analysis.

The Tn-Seq analysis was based on previous work (33). Genomic DNA was isolated from pooled samples using the Qiagen Tissue gDNA kit. gDNA was then cut with MmeI and DNA corresponding to the size of the transposon was purified by gel extraction. Adapters containing unique barcodes and priming sites for Illumina sequencing were ligated onto the DNA ends. The unique transposon insertion sites were amplified by PCR using primers specific to the adapter and the transposon, then purified by PCR cleanup kit and quantified. The DNA was sequenced using a HiSeq 2500 machine by the TCAG DNA Sequencing Facility. Results were analyzed using Qiagen’s CLC Genomics software using EDGE testing to generate fold-change and significance scores comparing Tn-library prey recovered from active killer strains and the corresponding *ΔtssM* control. Genes of interest were selected for follow-up analysis based on a p-value < 0.05 and a fold-change greater than 75-fold for *V. cholerae* prey or 10-fold for *E. coli*. The gene sets were then trimmed manually to test the most promising candidates for novel findings by removing genes known to influence T6SS survival (ex. immunity genes), tRNAs, and redundant genes.

### Statistical analyses

EDGE test analyses of Tn-Seq data were conducted using the CRC Genomics software. Other statistical analyses were computed using GraphPad Prism.

## Acknowledgements

This work was supported by grants from National Key R&D Program of China (2020YFA0907200), National Natural Science Foundation of China (31770082 and 32030001), Canadian Institutes of Health Research (CIHR) and Canadian Natural Sciences and Engineering Research Council (NSERC) to T.G.D.. T.G.D. was also supported by a Government of Canada Research Chair award, and a Canadian Foundation for Innovation grant (CFI-JELF). S.J.H. was supported by a CIHR Postdoctoral Fellowship for this work and K.M. by an Alberta Queen Elizabeth II Graduate Scholarship.

## Author contributions

S.J.H. and T.G.D. conceived the project; S.J.H., R.T.S., and K.M. designed, performed and analysed experiments; S.J.H. prepared the original manuscript and figures. T.G.D. supervised the study and guided manuscript revision.

## Supplemental material legends

**Figure S1:**
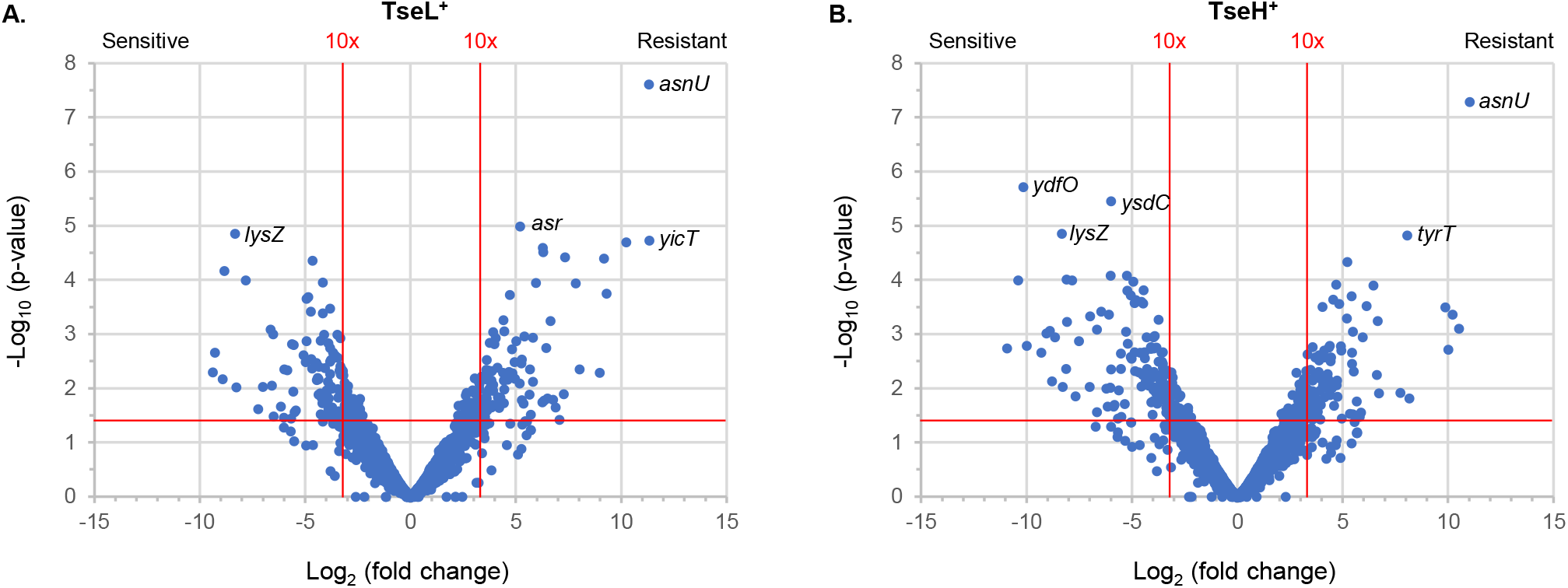
*E. coli::Tn* survival against *V. cholerae* TseL^+^ or TseH^+^. **A.** Volcano plot overviewing Tn-Seq data with *E. coli* prey. X-axis shows fold change comparing *V. cholerae* killer strains with only TseL intact (TseL^+^) or all antibacterial effectors inactivated (4eff_C_). A negative number shows reduced abundance in the sample treated with TseL^+^.Vertical red lines indicate ~10-fold changes. Y-axis shows p-value of EDGE test statistical analysis. Horizontal red line indicates ~0.05. **B.** As in **A** but showing data for TseH^+^ killers.

**Figure S2:**
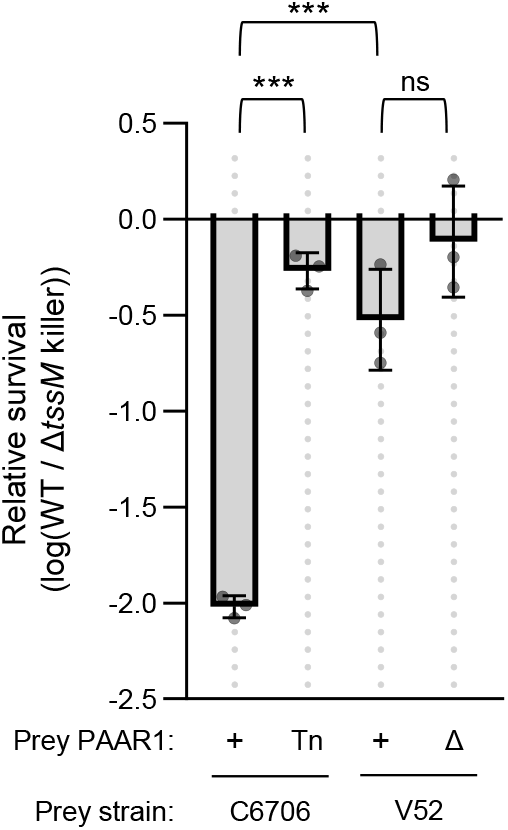
PAAR1 mutant survival in *V. cholerae* C6706 compared to V52. Log CFU recovered for stationary phase *V. cholerae* prey strains after competition with log-phase V52 killers. Prey strains have intact PAAR1 (+), PAAR1 deleted by scarless homologous recombination (Δ), or a transposon insertion in PAAR1 (Tn). Data show (as log) the ratio of prey CFU recovered after incubation with wild-type (WT) / Δ*tssM* killers. One way ANOVA with Tukey’s multiple comparisons tests: *** p < 0.001; ns, not significant.

**Figure S3:**
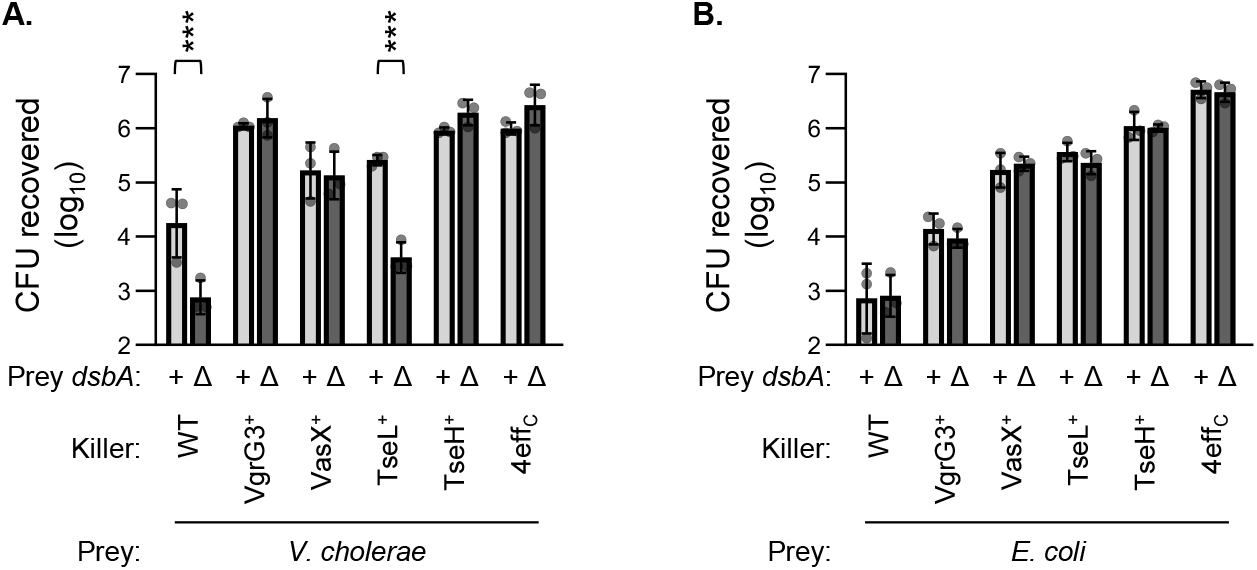
CFU recovery data relating to Figure 4. **A, B.** Log CFU recovered for *V. cholerae* (**A**) or *E. coli* (**B**) prey with their *dsbA* gene intact (light bars, ‘+’) or interrupted (dark bars, ‘Δ’). Killer strains were wild-type (WT) *V. cholerae*, a strain with all antibacterial effectors catalytically inactivated (4eff_C_), or strains with only one active effector (indicated). One way ANOVA with Sidak’s multiple comparisons test comparing prey with and without active *dsbA*: *** p < 0.001.

**Supplemental Table S1: Tn-Seq dataset**

**Supplemental Table S2: Strains and plasmids used in this study**

